# A correlative study of the genomic underpinning of virulence traits and drug tolerance of *Candida auris*

**DOI:** 10.1101/2023.04.07.536049

**Authors:** Bo Yang, Benjamin Vaisvil, Daniel Schmitt, Joseph Collins, Eric Young, Vinayak Kapatral, Reeta Rao

## Abstract

*Candida auris* is an opportunistic fungal pathogen with high mortality rates that presents a clear and present threat to public health. The risk of *C. auris* infection is high because it can colonize the body, resist antifungal treatment, and evade the immune system. The genetic mechanisms for these traits are not well-known. Identifying them could lead to new targets for new treatments. To this end, we present an analysis of the genetics and gene expression patterns of *C. auris* carbon metabolism, drug resistance, and macrophage interaction. We chose to study two *C. auris* isolates simultaneously, one drug sensitive (B11220 from Clade II) and one drug resistant (B11221 from Clade III). Comparing the genomes, we found that B11220 was missing a 12.8 kb gene cluster encoding proteins related to alternative sugar utilization, possibly L-rhamnose. We show that B11221, which has the cluster, more readily assimilates and utilizes D-galactose and L-rhamnose. B11221 also exhibits increased adherence and drug resistance compared to B11220 when grown in these sugars. Transcriptomic analysis of both strains grown on glucose or galactose showed that genes associated with translation were upregulated in B11221, and the putative L-rhamnose gene cluster was upregulated when grown on D-galactose. These findings reinforce the growing evidence of a link between metabolism and tolerance. Since immune system evasion depends heavily on the cell surface, we characterized cell wall composition and macrophage evasion for the two strains. We found that B11221 has far less β-1,3-glucan exposure, a key determinant of immune system evasion, and resists phagocytosis by macrophages compared to B11220. In a transcriptomic analysis of both strains co-cultured with macrophages we found that B11221 upregulates genes associated with early stages of growth and transcription factors that regulate transport. These key differences in growth and membrane composition could explain the resistance to phagocytosis and increased stress tolerance in general of B11221, and indicates another connection between metabolism and immune system evasion. Taken together, these data show that membrane composition, metabolism, and transport all correlate with colonization, drug resistance, and immune system evasion in *C. auris*.

## 1 Introduction

*Candida auris* was first clinically isolated in 2009 and has been rapidly reported in the years since in over 30 countries worldwide [1, 2]. Thus, it has emerged as a public health threat in the last decade, officially categorized as an urgent threat level by the Centers for Disease Control [3]. Furthermore, the World Health Organization has noted that *C. auris* causes hospitalization and death and urges more research and drugs to treat this pathogen [4]. The yeast can cause severe systemic infections, especially in immune-compromised individuals, with mortality rate of greater than 40% due to limited treatment options [5–7]. Effective *C. auris* infection treatment is challenging because it exhibits multi-drug resistance to antifungals [8], it persists in hospital environments with transmission within health care settings [8–10], and misidentifications delay diagnosis [11, 12]. From work on other *Candida* pathogens, it is known that virulence is a function of metabolic flexibility, host adherence, biofilm formation, filamentation and secretion of hydrolytic enzymes [13]. These virulence factors and fitness attributes of *C. auris* are not well-known. Therefore, we undertook a genomic and transcriptomic study of these attributes, comparing two different *C. auris* isolates under different growth conditions with non-canonical carbon sources, antifungal treatments, and macrophages.

1. *C. auris* has been placed within the *Clavispora*/*Candida* clade of the Metschnikowiaceae family of the order *Saccharomycetales*, yeasts that reproduce by budding [14]. *C. auris* is in the CTG clade, a group of yeasts that translate the codon CTG to serine instead of leucine [15], although it is genetically distant from the other widely studied CTG clade species including *C. albicans*, *C. tropicalis*, and *C. parapsilosis* [16]. Previous studies suggested that *C. auris* emerged simultaneously and independently in different global regions [17]. Based on a variety of criteria such as genome sequence, geographic location of isolation and drug resistance profile, *C. auris* isolates have been classified into five clades by geographic regions: Clade I (South Asian), Clade II (East Asian), Clade III (South African), Clade IV (South American) [18], and Clade V (Iran) [19]. The first four clades are separated by thousands of single-nucleotide polymorphisms (SNPs), although within each clade the isolates appear to be clonal [17]. Clade V (Iran) is most recently discovered, and is also separated from the other clades by >200,000 SNPs [19]. Genome assemblies first came from Clade I isolates, such as Ci6684 [20] and five isolates from an outbreak in United Kingdom [21]. Muñoz et al. [16] assembled and compared genomes across Clade I to IV. *C. auris* genomes have been mapped into seven chromosomes using optical maps, which were then confirmed by long-reads and telomere-to-telomere assemblies [22]. One isolate from Iran representing Clade V has been sequenced so far and its genome is highly syntenic with those of Clades I, III, and IV, which are the outbreak causing clades [19, 22].

The CDC reports that 90% of *C. auris* isolates are resistant to at least one antifungal and 30% are resistant to at least two antifungals [23], making them multidrug resistant. Antifungals currently in clinical use fall in four major classes: azoles, polyenes, allylamines and echinocandins [24]. The first three classes inhibit ergosterol [25], an essential component of the fungal cell membrane, while echinocandins inhibit β-1,3-glucan [26], an essential component of the fungal cell wall. Both ergosterol and β-1,3-glucan are not present in mammals. Susceptibility testing has shown that over 90% of the isolates are resistant to azoles, 10-35% are resistant to the polyene amphotericin B, yet only 4-7% are resistant to echinocandins [17, 27, 28]. Another report stated that 44% of *C. auris* isolates are resistant to the azole fluconazole, 15% to amphotericin B, yet only 3% are resistant to the echinocandin caspofungin [29]. Understanding of the molecular mechanisms of the drug resistance in *C. auris* is limited. However, studies have correlated azole and polyene resistance to mutations in the ergosterol pathway, specifically *ERG11*, *ERG2, ERG3,* and *ERG6* [30] [14, 31–33]. Other mechanisms have been proposed, like variation in *ERG11* copy number, overexpression of efflux pumps, and changes in membrane composition that protect the target [34, 35]. While only a small number of strains are resistant to echinocandins, *C. auris* has an *FKS1* gene that encodes for β-1,3-d-glucan synthase that shares the two hot spots for resistance mutations in *Candida spp.* [36] [37–40] [41]. Taken together, these results point to mutations in ergosterol biosynthesis as the primary reason for multidrug resistance in *C. auris*, and highlight β-1,3-glucan biosynthesis as the most effective target pathway. As a result, echinocandins have become the drugs of choice for primary treatment of invasive *C. auris* infections [42].

Antifungal drug tolerance has been proposed to increase the frequency of drug resistance [43]. Yet, *C. auris* drug tolerance has not yet been investigated. Drug tolerance is defined as the ability of a subpopulation of drug-susceptible fungi to grow slowly in a drug above the MIC [44]. In the laboratory, tolerance is quantified using disk diffusion assays or broth microdilution assays after 48 h of growth [44]. In disk diffusion assays on solid media, drug tolerant cells can be visualized as they grow inside of the zone of inhibition. This manifests clinically as a sub-population of patients with persistent infections that recur after treatment[45]. Thus, distinguishing drug tolerance from resistance is important to provide insights to treatment failure. The genetic underpinning of drug tolerance in yeasts is not well-known, although point mutations, aneuploidy, loss of heteroresistance, growth conditions, stress response pathways, and mutations in *TAC1*, a positive transcriptional regulator of efflux pumps, have all been implicated [46] [47].

While drug resistance and tolerance affect treatment, carbon source utilization appears to be a key determinant of primary infection. The carbon utilization profile of *C. auris* has not been well-established. However, adaptation of *Candida* spp. to different carbon sources has been connected to virulence, primarily because it permits colonization of glucose-limited niches. *Candida* species assimilate local nutrients to counter local environmental stresses and evade local host defenses [48], which impacts the outcomes of fungal pathogen infections [49]. Compared to glucose, *C. albicans* grown on lactate has a thinner cell wall with reduced β-1,3-glucan and chitin, enhancing antifungal resistance [50]. Furthermore, they are better at evading the immune system – cells are less likely to be taken up by macrophages, and are more efficient at killing and escaping them [51]. Acetate-grown *C. glabrata* are more susceptible to fluconazole, generate less robust biofilms, and are more susceptible to macrophage killing, possibly due to upregulation of carboxylate transporters and a potential role of *FPS1* and *ADY2a* in the phagocytosis process [52].

Interaction between a microbe and the host is mediated through the recognition of pathogen-associated molecular patterns (PAMPs) by pattern recognition receptors (PRRs) on immune cells. PAMPs derived from fungal pathogens are specifically recognized by PRRs, occurring at the interface between the immune cell membrane and the fungal cell wall. The major fungal PAMPs on the fungal cell surface include β-glucan, mannan and chitin, which are critical components of the fungal cell wall [53]. These have not been well-characterized in *C. auris.* However, polysaccharide fibrils of β-(1,3)-glucan covalently linked to β-(1,6)-glucan and chitin compose the core structure of *Candida* spp. cell wall and function as a scaffold for external proteins [54]. β-glucan layers lie between chitin and mannan layers and trigger strong strong inflammatory responses [55], however, it is normally masked by the less pro-inflammatory mannan layer but can be exposed during infection [56, 57]. β-glucan recognition by its receptor, dectin-1, is required to control systemic infections [58] and plays a major role in anti-*Candida* immune responses [53]. Its biosynthesis is critically dependent on β-(1,3)-glucan synthase activity (encoded by *FKS1* and *FKS2* genes [59]), which is the target of echinocandins such as caspofungin.

In this study, we chose two *C. auris* strains that have different antifungal resistance traits. They are both from the most prevalent *C. auris* clades with many cases reported and wide geographical distribution [60]. *C. auris* B11220 (aka B11220^Δ^) is from Clade II (East Asia), reported with ear infection cases, susceptible to all antifungal drugs and not associated with outbreaks; *C. auris* B11221 is from Clade III (South Africa), reported in outbreaks of invasive and multidrug-resistant infections. *C. auris* B11221 is more drug resistant, the MIC-50 (Minimum Inhibitory Concentration required to inhibit the growth of 50 percent) to caspofungin 16 μg/mL for B11221 and only 0.125 μg/mL for B11220, according to the CDC.

Comparing the two *C. auris* genomes showed that a gene cluster putatively encoding an L-rhamnose utilization cluster was missing in B11220 (but present in B11221, which is more drug resistant and has a higher immune cell survival than B11220^Δ^. To indicate this, we denoted the deletion strain B11220^Δ^. In alternative carbon sources, B11221 utilized sugars more readily, reduced abiotic surface adhesion, and developed drug tolerance. We found upregulated genes in D-galactose grown B11221 are associated with the ribosomal complex, translation factors, protein metabolism and in carbohydrate metabolism. Several ORFs unique to B11221 were also found to function in stress response, cell wall biosynthesis and carbohydrate metabolism. Transcriptomic analysis of the interaction of both strains with macrophage discovered up-regulation of transport-related transcription factors in B11221, which may contribute to its higher stress tolerance and better host survival.

## 2 Materials and methods

### 2.1 Candida growth

1. *C. auris* strains were stored at -80°C prior to use. Isolates from frozen stock were streaked out on yeast-peptone-dextrose (YPD) agar at 30°C for 24 hours and subsequently stored at 4°C prior to liquid culture. Single colony of each strain was picked and grown in 50ml YPD medium at 30°C at 200 rpm for 16hrs. Then cells were pelleted by centrifugation (3,000 × g) and washed three times in phosphate-buffered saline (PBS). The cells were then adjusted to the desired concentration after measurement with an optical spectrometer and then resuspended in selected media for each assay, as described in each result section.

For growth curve assays, each well of a flat-bottom 96 well polystyrene plate was loaded with 200 μL of an inoculum of 0.05 OD/mL in Synthetic Complete (SC) Medium. Different carbon sources were added to a final concentration of 2 %. Antifungal drugs were added as described. The experiments were conducted in triplicates. The plates were incubated at 30 °C for 45 hours and measured at 600 nm every 15 min in a BioTek Synergy H1 plate reader with shaking. Growth curves fitted using non-linear model in Graphpad.

### 2.2 Genome sequencing and analysis

#### Genome sequencing and Assembly

High-molecular weight genomic DNA was isolated based on a modified version of Promega’s Genomic DNA Isolation Kit (Promega, A1120) [63]. Nanopore libraries were prepared with the Rapid Barcoding Kit (ONT, SQK-RBK004). Illumina libraries were prepared with the Nextera DNA Flex Library Prep Kit (Illumina, 20018704) along with the Nextera DNA CD Indexes (Illumina, 20018707). Nanopore sequencing was performed with the Oxford Nanopore MinION and MinKNOW software, and the resulting fastq files were demultiplexed using EPI2ME (Metrichor, Oxford, UK). Illumina sequencing was performed with Local Run Manager software on the iSeq 100 machine. A GENERATEFASTQ run was initiated and run with the parameters Read Type: Paired End, Read Lengths: 151, and Index Reads: 2. Reads were demultiplexed using the native software on the iSeq 100 machine. The *C. auris* B11220 and B11221 sequences were assembled using the Prymetime (v0.2) pipeline [61], which is a hybrid assembler using long and short reads.

#### Data availability

Data for *C. auris* B11220 and *C. auris* B11221 are available on NCBI under BioProject PRJNA865346. *C. auris* B11220 has been assigned the NCBI BioSample accession SAMN30106616. The whole genome assembly can be accessed with accession JANQCM000000000. The raw reads are available at the NCBI Sequence Read Archive (SRA) – nanopore reads are available under accession SRR20993210 and Illumina reads are available under accession SRR20993211. *C. auris* B11221 has been assigned the NCBI BioSample accession SAMN30106621. The whole genome assembly can be accessed with accession JANQCN000000000. The raw reads are available at the NCBI Sequence Read Archive (SRA) – nanopore reads are available under accession SRR20993402 and Illumina reads are available under accession SRR20993403.

#### Sequence analysis

The genomes of two *C. auris* strains were aligned to each other. Genome alignment was performed with ERGO’s Genome Align tool which uses BLASTN [62] with an expected threshold of 1E-180 to identify blocks of homology, which are then visually inspected on a genome browser. ORF functions were determined via protein similarity to orthologs with functions using the ERGO database as detailed in Overbeek et al. [63]. Additionally, domain analysis was performed using the NCBI Conserved Domain Database [64]. Annotations were then verified through manual curation.

### 2.3 Carbohydrates assay

Total Carbohydrate Assay Kit (Sigma Catalog Number MAK104) was used for this assay. PBS-washed overnight cultures of *C. auris* strains were inoculated to SC media respectively with 2% D-glucose, 2% D-galactose and 2% L-rhamnose, at concentration of 0.1 OD600. Then the cultures were distributed to 96 well plate at 40 μL per well. Media without *Candida* cells were used as controls. Standard curves were created using 2 mg/mL standard solutions. All conditions were in triplicate, incubated at 30°C for 24 h. Reaction assays were followed as described by the protocol provided in the Kit. Colorimetric detection was taken at the end of assay at absorbance of 490 nm (A490) to measure carbohydrates left in the media after *Candida* growth. Carbohydrates consumption by growth was calculated by subtracting the remaining sugar from before growth.

### 2.4 Surface adhesion assay

PBS-washed overnight cultures of *C. auris* strains were inoculated to SC media respectively with 2% D-glucose, 2% D-galactose and 2% L-rhamnose, at concentration of 0.5 OD600. Cell suspension (200 μL) was aliquoted into each well of a flat-bottom 96 well polystyrene plate. Each condition was repeated in triplicate. The plates were covered and incubated at 37°C for 4 hours. After incubation, unattached cells were discarded and 40 μL of 0.5 % Crystal Violet (CV) solution was added to each well for 45 minutes at room temperature to stain the attached *Candida* cells to the polystyrene plates. The excess of dye was discarded, and plates were washed 6-times with diH2O and gently tapped onto a paper towel to remove residual diH2O. To dissolve the CV in the attached cells, 200 μL of 75% methanol was added to each well and the plates were incubated at room temperature for 30 minutes. Absorbance of adhesion-retained CV dye was measured at 590 nm in Victor3 plate reader (PerkinElmer) and used as a measurement of adhesion to plastic surface.

### 2.5 Caspofungin susceptibility by disk diffusion assay

Overnight cultures of *C. auris* strains were washed in PBS and were standardized to 0.5 OD600 in PBS. About 600 uL was placed on each SC agar respectively containing 2% D-glucose, 2% D-galactose and 2% L-rhamnose in petri dishes. L-shape spreader was immediately used to spread the culture evenly on agar. After drying plates for one hour, one paper disk saturated with caspofungin solution (4 mg/mL in DMSO, 5 uL) was placed in center of each plate. These plates were kept in 30°C bottom-up. The zone of growth inhibition surrounding the filters (halo) was observed and photographed after 24 to 72 hours of incubation. Each condition was repeated in triplicates.

### 2.6 Cell survival by *Ex vivo* macrophage killing assay

Murine macrophages cell lines were derived from the mouse line NR-9456 and are available through BEI resources. Macrophage cells were thawed and grown in Dulbecco’s Modified Eagle Medium (DMEM) plus 10% fetal bovine serum (FBS) to reach the 95% confluence. *Candida* cells were grown in YPD liquid media at 30°C overnight prior to the infection experiment, then acclimated to macrophage media (DMEM+10%FBS) at 37 °C for one hour prior to the infection. On the day of the infection experiment, macrophages were seeded in 24-well plates at density of 0.5 ×10^6^ cells/mL and incubated at 37 °C for 4 hours for cells to attach to plate bottom. The cells were counted by hemocytometer and seeded to each well in a ratio of 1 *Candida* cell to 15 macrophage cells. The mixture plates were then incubated at 37 °C (5% CO_2_) for four hours and eight hours. The cultures were then scraped down from the wells using cell scrapers. The contents from each well were transferred to the collection tubes with 0.02% Triton X-100 to osmotically lyse the macrophages. A serial dilution was quickly performed to make the suspension diluted to 10^-4^. 150 µl of the dilution were spread on YPD agar plates and incubated at 30° C for 24 hours to make colony forming units (CFU). *Candida* survival was calculated by taking the ratio of CFU obtained from *Candida* and macrophage mixture to that obtained from *Candida* alone condition. Each condition was repeated in triplicates.

### 2.7 Quantification of β-1,3-glucan

#### Total cellular levels of β-1,3-glucan

*Candida* strains were grown overnight and diluted to an OD600 to1. Then the diluted cultures were killed by incubation in heat block at 100° C for 5 min. 100 μL of each strain was added to 1.5ml EP tubes. Tubes were centrifuged for 5 min at 3000×g. Harvested cells were resuspended in 50 μL PBS containing 2% BSA and incubated at room temperature for one hour using a swinging mixer. Tubes were centrifuged for 5 min at 3000×g and cells were resuspended in 50 μL of the antibody solution (antibody against β-1,3-glucan, Biosupplies, Cat. No. 400-2), then incubated at 37° for two hours on a swinging mixer. Tubes were centrifuged for 3 min at 3000×g and wells were washed three times with PBS containing 0.05% Tween-20 (PBS-T). Cells were then resuspended in 50 μL of the secondary antibody solution (1 μg/ml Abcam rabbit polyclonal anti-mouse 97046, 1:1000). Tubes were incubated at 37° for one hour on a swinging mixer/belly dancer. These tubes were centrifuged for 5 min at 3000×g and were washed three times with PBS-T and finally resuspended in 50 µl/well chemiluminescent peroxidase substrate (Supersignal^®^ West Pico ThermoFisher Cat No. 34077). The samples were then transferred to a reading plate, and luminescence was measured in a Victor3 plate reader (PerkinElmer).

#### β-1, 3-glucan surface exposure

*Candida* strains were grown overnight, washed, and diluted to an OD600 to 0.6. Each sample was blocked with 3% BSA in PBS then stained anti-β (1,3)-glucan antibody (Biosupplies Australia Pty Ltd., Australia) followed by secondary antibody of anti-Mouse IgG (H+L) Alexa Fluor 488 (ThermoFisher). The level of β-1,3-glucan exposure on the cell surface was then quantified on a Beckman Coulter CytoFlex 3-laser cytometers. Controls included single-stained and unstained samples for each strain. Flow cytometry data were analyzed using the Flowjo software to calculate the relative amounts of cell wall β-1,3-glucan.

### 2.8 Transcriptional profiling by RNA-sequencing

#### Sample preparation

##### Carbon source study

Overnight cultures of *Candida* strains were washed three times by PBS. 15 OD600 of each strain were inoculated to SC media respectively containing 2% D-glucose and 2% D-galactose, in three replicates. All cultures were incubated at 30° C while shaking at 220rpm for 4 hrs. Then the cultures were spun down to remove supernatant and pellets were immediately transferred to liquid nitrogen and stored in -80 °C.

##### Host-pathogen interaction

Assay was followed similarly as described in Methods 2.6 *Ex vivo* macrophage killing assay for *Candida* survival. Modification was made to mix each *Candida* strain and macrophages cells to each well in a ratio of 1:1 *Candida* cell to macrophage cells in DMEM containing 10% FBS. The mixture cultures in 24-well plates were incubated at 37 °C (5% CO_2_) for four hours. The cultures were then scraped down from the wells using cell scrapper. The contents from each well were transferred to collection tubes then centrifuged to remove supernatant. Cell pellets were immediately transferred to liquid nitrogen and stored in the -80 °C freezer.

##### RNA sequencing

RNA was extracted using the Qiagen RNeasy kit following the manufacturer’s directions. PolyA mRNA was used as input to SMARTer Stranded RNA-Seq Kit (Takara Biosystems) according to manufacturer’s instructions for library preparation. The library was sequenced on Illumina HiSeq 2×150 obtaining approximately 593 million reads. The sequenced reads were checked for quality using ERGO’s (Overbeek et al., 2003) read QC workflow. Reads were then quantified using kallisto (Bray et al., 2016; version 0.46.1) through ERGO’s RNA-Seq workflow. The transcript abundances were summarized by tx2gene R package (Soneson et al., 2015; version 1.20.0) and subsequently converted into counts. Gene counts were imported into DESeq2 R package (Love et al., 2014; version 1.16.1) and tested for differential expression using DESeq2’s deseq function. Genes were considered significantly differentially with a log2 fold change greater than 2 and a false discovery rate (FDR) adjusted p value less than 0.5. Genes identified as differentially expressed in all carbon sources were filtered.

##### Data availability

RNA sequencing data for *C. auris* B11220 and *C. auris* B11221 are available at the NCBI. Accession for Sequence Read Archive (SRA) data is PRJNA902676, submission ID: SUB12286496.

## 3 Results

### 3.1 A 12.8 kb deletion region encodes genes involved in alternate sugar assimilation

We first compared the genomes of the *C. auris* isolates B11220 and B11221, which not only differ in drug susceptibility, but originate from distant geographic regions as well. B11220 is from Clade II (East Asia) and sensitive to the three major classes of antifungal drugs, while B11221, from Clade III (South Africa) is resistant to these drugs. These isolates have previously been sequenced [22]. We resequenced these isolates using the Prymetime (v0.2) pipeline which uses both long reads and short reads to obtain a more contiguous genome assembly [61]. Using ERGO’s Genome Align tool, we confirmed that there is a 12.8 kb region absent in B11220 but present in B11221. Looking into the other available isolate genome sequences, we found that it is absent in Clade I, II, IV isolates and present in Clade III and Clade V [22]. This region contains seven predicted open reading frames (ORFs). This region contains seven predicted open reading frames (ORFs) that code for transport and degradation of alternate sugars in general but the closest homologs are genes involved in L-rhamnose (Fig. 1) metabolism from *Aspergillus* [65–67]. These ORFs are identified as (RTCAU02308) *LRA1* (LraA-L-rhamnose 1-dehydrogenase, EC 1.1.1.173), (RTCAU02624) *LRA2* (LrlA-l-rhamnono-γ-lactonase, EC 3.1.1.65), *LRA4* (LkaA-2-keto-3-deoxy-L-rhamnonate aldolase, EC 4.1.2.53), (RTCAU02415) major facilitator superfamily (MFS) sugar transporter, (RTCAU02487) *ramA* (rgxB-Alpha-L-rhamnosidase, EC 3.2.1.40), (RTCAU02564) *RhaR* (transcriptional regulator involved in the release and catabolism of the methyl-pentose [68]) and (RTCAU02624) *LRA3* (LrdA l-rhamnonate dehydratase, EC 4.2.1.90). ORF functions were determined via protein similarity to orthologs with functions using the ERGO database as detailed in Overbeek, et al [63]. Additionally, domain analysis was performed using the NCBI Conserved Domain Database [64]. Annotations were then verified through manual curation. For reader convenience, we will identify the isolate with the deletion with a delta superscript (B11220^Δ^) to distinguish it from B11221 which retains the seven open reading frames (ORFs). We hypothesize that this large genomic difference could be linked to phenotypic variation of *C. auris*. Here we demonstrate a connection between these seven genes with respect to carbon assimilation and drug tolerance of *C. auris*.

**Figure 1.**
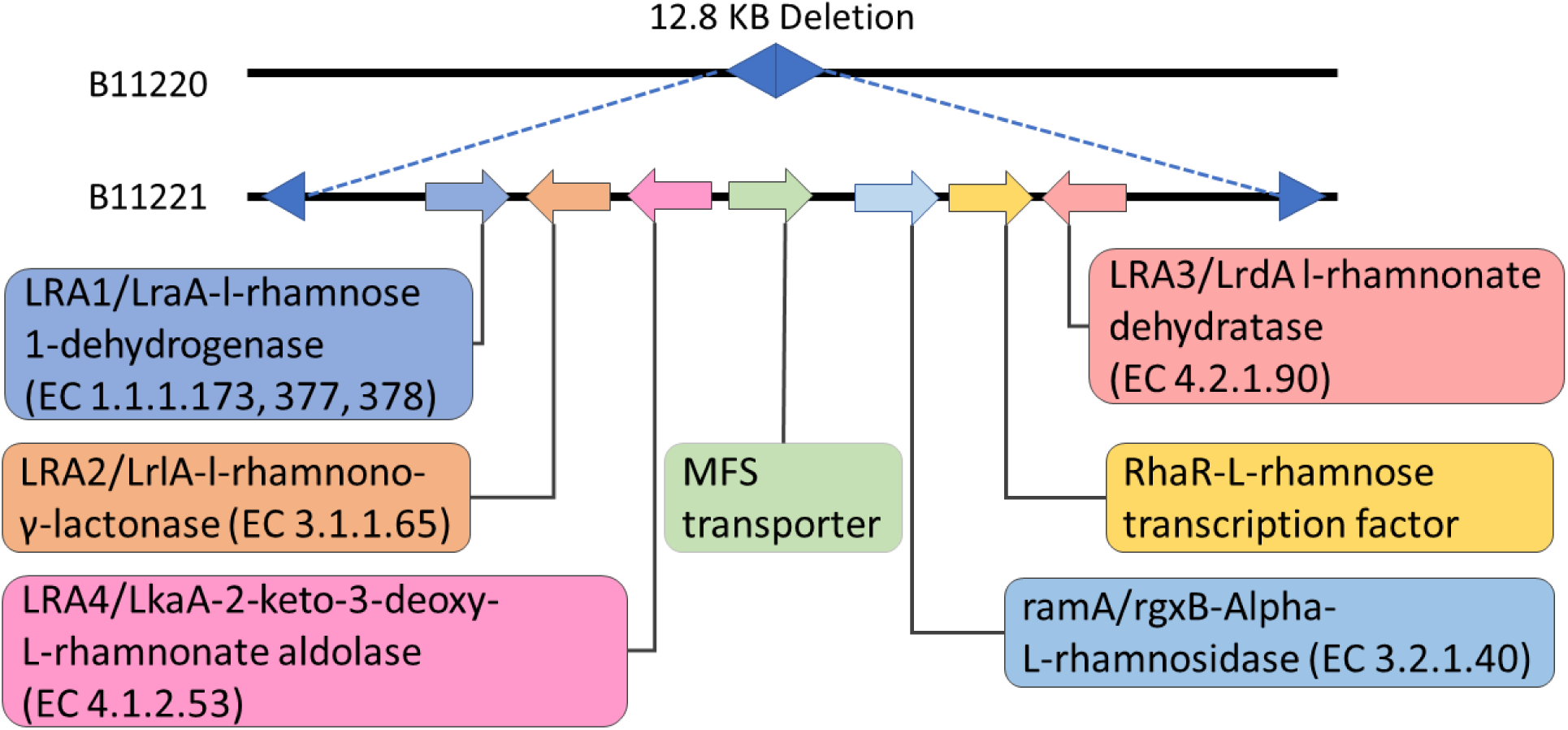
12.8 kb region absent from B11220^Δ^ compared to B11221. Location and regions of the seven closest ORFs for L-rhamnose digestion. Black lines are homologous between the two genomes.

L-rhamnose (l-6-deoxy-mannose) is a C6 sugar enriched in some plant fractions like hemicellulose and pectin. It can be used as a carbon and energy source by several microorganisms [69]. There are two pathways known for L-Rha catabolism: one with phosphorylated intermediates described in bacteria, and the other one without phosphorylated intermediates described in yeast species like *Pullularia pullulans* [70], *Pichia stipitis* and *Debaryomyces polymorphus* [71]. The enzymes in the fungal L-Rha catabolism pathway include l-rhamnose-1-dehydrogenase (EC 1.1.1.173), l-rhamnono-1,4-lactonase (EC 3.1.1.65), l-rhamnonate dehydratase (EC 4.2.1.90), and l-erythro-3,6-dideoxyhexulosonate aldolase (EC 4.1.2.) [69]. These genes are involved in L-Rha transport from extracellular space for a four enzymatic steps conversion to a final pyruvate compound for carbohydrate metabolism [72].

Although the seven deleted genes were first described as being involved in L-rhamnose metabolism in yeast and bacterial species, we hypothesized that they are likely promiscuous and involved in the metabolism of non-canonical sugar sources, since many proteins involved in transport and catabolism of alternative sugars are promiscuous [73]. To find if the missing ORFs affect alternative sugar metabolism for *C. auris*, we performed spot assays for D-glucose (control), L-arabinose, D-galactose [29], α-lactose, D-maltose, L-rhamnose, sucrose and D-xylose (Fig. S1). No difference between the two isolates was observed, so we further characterized growth on L-rhamnose and D-galactose with more quantitative methods. We used multiple readouts including plate reader growth curves, growth in liquid cultures, growth on solid agar media, and carbon source assimilation (Fig. 2). Our results showed that there was no significant difference in the plate reader growth curves between B11220^Δ^ and B11221 in D-galactose or L-rhamnose (Fig. 2a). When grown in the presence of D-glucose, B11220^Δ^ accumulated to a higher OD_600_ than B11221 in liquid media (Fig. 2b). Overnight liquid culture of B11221 was clumpy in D-glucose but the aggregative phenotype was not observed in D-galactose or L-rhamnose, while B11220^Δ^ cultures grew planktonically without aggregates in three sugars conditions (Fig. 2c). Small colonies as well as decreased number of cell colonies were observed in both B11220^Δ^ and B11221 when in D-galactose and L-rhamnose as the carbon source as compared to D-glucose (Fig. 2d). Strain B11220^Δ^ formed fewer colonies forming units (CFU) as compared to B11221 in each sugar at OD600 dilution of 0.001 and 0.0001 (Fig. 2d). Assimilation of D-glucose as well as D-galactose was indistinguishable between the two isolates, but L-rhamnose assimilation was significantly different (Fig. 2e). These results draw a positive correlation between the missing genes with L-rhamnose assimilation in *C. auris*, although there must be another pathway present that allows B11220^Δ^ to grow on L-rhamnose. Thus, we hypothesize that this cluster enhances, but is not necessary, for L-rhamnose catabolism in *C. auris*.

**Figure 2.**
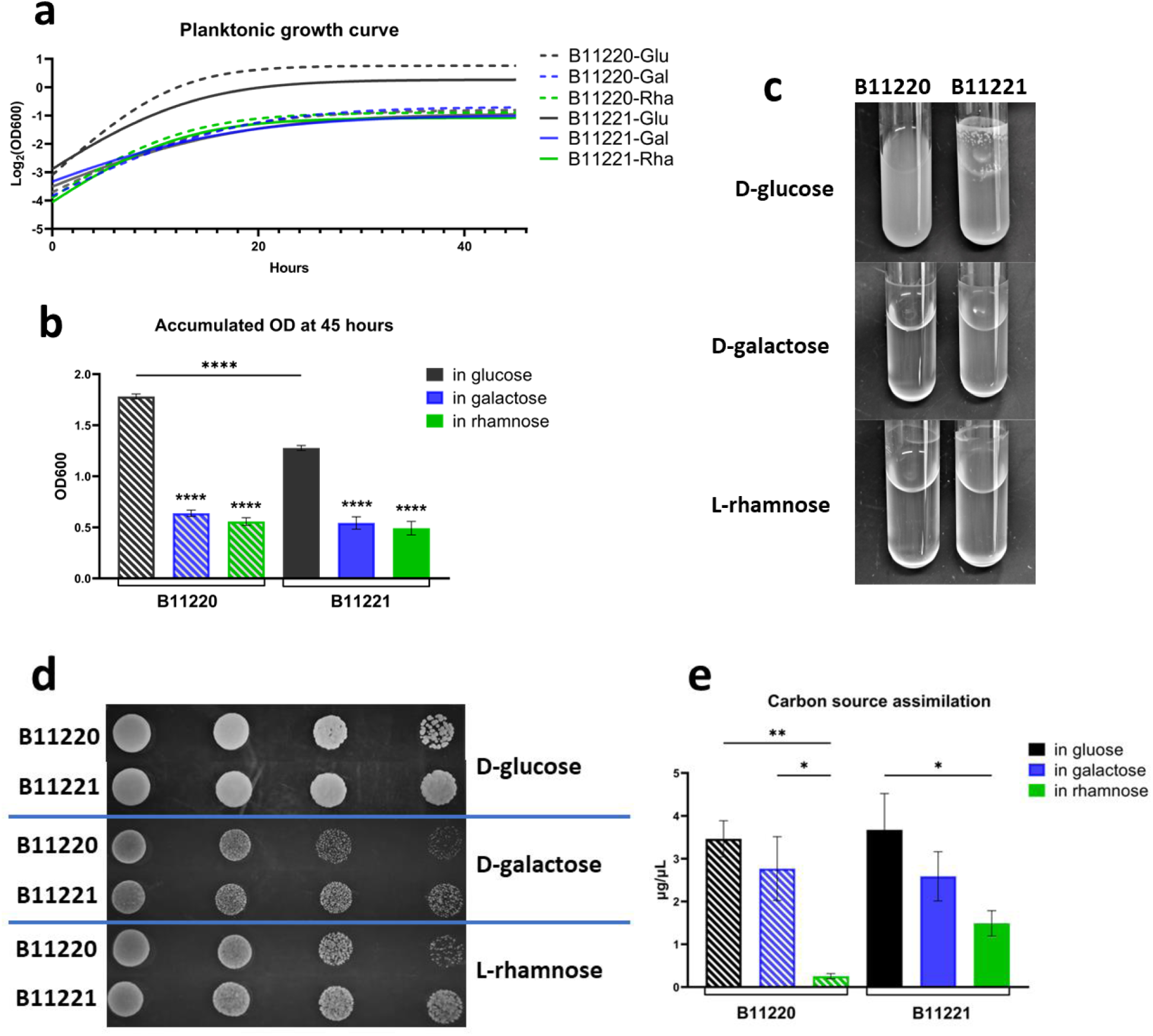
Growth of two *C. auris* strains in 3 sugars. (a) Planktonic growth of B11220^Δ^ (blue) and B11221 (green) in liquid SC media. Curve fit with non-linear model. Logged. (b) End point OD600 comparison of growth in liquid culture tubes at 45 hours. Underlined asterisk marks indicate significant differences of same sugar type between two strains, and asterisk marks without underline indicate significant differences between different sugars within one strain (****p<0.0001). (c) Liquid culture overnight grown in SC media with glucose, galactose and rhamnose in glass tubes. (d) Serial diluted colonies growth of B11220^Δ^ (top rows) and B11221 (bottom rows) on SC agar in 3 sugars. OD600 of colonies from left to right are 0.1, 0.01,0.001, 0.0001. (e) Assimilation of three sugars as carbon sources by two *C. auris* strains using Total Carbohydrate Assay Kit from Sigma. Tukey’s multiple comparisons test (****p<0.0001, **p<0.005, *p<0.05)

### 3.2 Alternative carbon sources increase drug tolerance of *C. auris* strain B11221

Alternative carbon sources can modulate the sensitivity of *Candida* species to antifungal drugs [48]. Therefore, we measured the drug sensitivity of *C. auris* B11220^Δ^ and B11221 when grown in D-glucose, D-galactose, and L-rhamnose. We exposed the microbes to a representative of each of the three classes of antifungal agents, fluconazole (azole), amphotericin B (polyene), and caspofungin (echinocandin). We tested the MICs of the two strains with the E-test assay and determined the MIC values (in μg/mL) of fluconazole to be 8.5 for B11220^Δ^ versus 128 for B11221; of amphotericin B to be 0.28 for B11220^Δ^ versus 0.35 for B11221; of caspofungin to be 0.028 for B11220^Δ^ versus 0.75 for B11221. These values are in line with CDC reports [74]. Based on this information, we used fluconazole at 10 μg/mL for B11220^Δ^ and at 130 μg/mL for B11221; Amphotericin B at 0.38 μg/mL for both strains; and Caspofungin at 0.2 μg/mL for B11220^Δ^ and at 16 μg/mL for B11221.

As expected, growth of *C. auris* B11220^Δ^ and B11221 is affected by antifungals with every carbon source (Fig. 3a). Both strains are equally inhibited by amphotericin B. However, B11220^Δ^ is particularly inhibited in fluconazole and does not grow in the presence of caspofungin, while *C. auris* B11221 is most inhibited by caspofungin but still grows (Fig. 3a). This is consistent with reports of B11221 being sensitive but tolerant to caspofungin [75].

**Figure 3.**
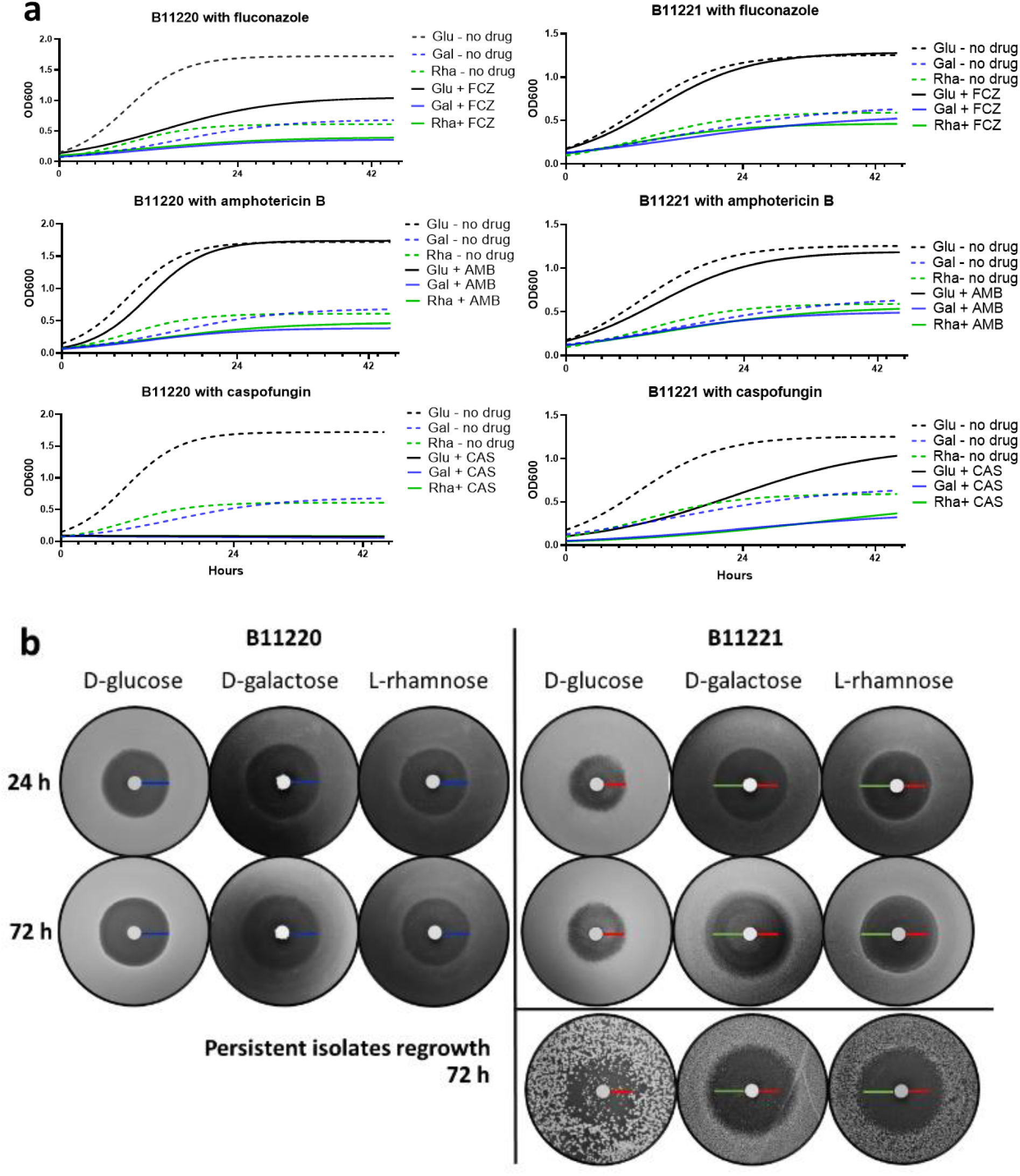
*C. auris* strains growth in the present of three antifungal drugs in three carbon sources. (a) Planktonic growth of two strains in liquid SC media. Curve fitting with non-linear model. Left panels: curves of B11220^Δ^. Right panels: curves of B11221. Top: in fluconazole, drug concentration was 10 μg/mL for B11220^Δ^, 130 μg/mL for B11221. Middle: in amphotericin B, drug concentration was 0.38 μg/mL for both strains. Bottom: in caspofungin, drug concentration was 0.2 μg/mL for B11220^Δ^, 16 μg/mL for B11221. Red line indicates the duration at 24 hours. (b) Photographs from disk diffusion assay showing growth of two C. auris strains on SC agar in three carbon sources in presence of caspofungin disks at concentrations of 1M. Left panel: B11220^Δ^ growth on SC agar plate with caspofungin for 24 hours (top) and 72 hours (bottom). Right panel: B11221 growth on SC agar plate with caspofungin for 24 hours (top) and 72 hours (middle). Drug tolerant persistent isolates of B11221 were picked and grown again on SC agar plates in three sugars with caspofungin disks for 72 hours (bottom). Photos are representative for biological repeats of the experiment four times. Agar plates were incubated at 30 °C. Scale bars in blue measure 9.14 mm (0.36”), in red measure 7.11 mm (0.28”), in green measure 10.16 mm (0.4”).

To further investigate drug tolerance, we performed a disk diffusion assay for each isolate on all three carbon sources). Drug tolerance is shown by colonies within the zone of growth inhibition (or halo), which indicates slow growth at concentrations above the MIC, and is measured as fraction of growth (FoG) [44] [45]. In the assay, the clear zones formed by *C. auris* B11220^Δ^ in each of the three sugars were comparable in size and maintained the area from 24 hours to 72 hours during growth, as indicated by the blue scale bar (Fig. 3b, left panel). However for B11221, the halos formed in D-galactose and L-rhamnose were larger compared to that in D-glucose after 24 hours growth (Fig. 3b, right panel), suggesting a lower MIC to caspofungin in these alternative sugars [44]. This phenotype is consistent with the growth patterns we observed in aqueous media (Fig. 3a, bottom panel). 48 hours later, colonies appeared inside the zone of inhibition for *C. auris* strain B11221in all three sugars, indicating FoG above the MIC and therefore drug tolerance [44]. The drug tolerance phenotype was especially pronounced in D-galactose (Fig. 3b, right middle panel)

Drug tolerant B11221 colonies were then harvested and regrown on SC agar in the presence of each of the three sugars with or without caspofungin (Fig. 3b, right bottom panel). Compared with the first round of growth, D-glucose regrown-colonies were bigger in size, maintaining a zone of inhibition, and tolerant isolates appeared as early as 24 hours of incubation. On D-galactose and L-rhamnose plates, the second-round inhibition zones were similar in size to first-round suggesting their MIC to caspofungin remained. In addition to the zone area, there was no colony regrown when in L-rhamnose suggesting a loss of drug tolerance. Fewer tolerant colonies were able to develop within the inhibition zone in D-galactose, and these appeared to maintain a drug tolerance to caspofungin in presence of D-galactose. Together our results suggest *C. auris* B11221is tolerant to caspofungin especially in the presence of alternate sugars.

### 3.3 Alternate carbon sources reduce adhesion and aggregation of *C. auris* B11221

Adhesion and aggregation affect virulence via colonization, biofilm formation on surfaces, and ultimately dissemination to initiate infections at distal locations [76–80]. Studies have shown that *C. auris* can survive on both moist and dry surfaces for long periods of time and still be cultured [81, 82], and it more inclined to adhere and form biofilms on catheters compared to *C. albicans* [83]. The persistence of *C. auris* on abiotic surfaces presents opportunities for it to colonization and spread rapidly within health care facilities. We characterized the adhesion of B11220^Δ^ and B11221 to polystyrene and agar when grown on D-glucose, D-galactose, and L-rhamnose. The alternate sugars did not affect adhesion for B11220^Δ^ (Fig. 4), but B11221 adhered significantly less in D-galactose and L-rhamnose as compared to D-glucose (Fig. 4b). Furthermore, B11221 adhered better than B11220^Δ^ in all three carbon sources on both polystyrene (Fig. 4a and 4b) and agar (Fig. 4c). In addition to adhesion, the aggregative phenotype of B11221, which is linked to transcriptional changes in genes involved in cell surface adhesion [84], is also reduced in presence of non-canonical sugars (Fig. 2c). Looking into the genomes, we discovered that ORF (RTCAU00295), encoding cell-wall agglutinin, was unique to B11221 (Table 1) and not identified in B11220^Δ^. Agglutinin-like sequence proteins (Als) are cell surface glycoprotein of *Candida* species that play essential roles in the processes of adherence and biofilm formation *in vitro*. In our later transcriptomics experiment (see Section 3.4), we also observed that ORF was upregulated in D-glucose but not in D-galactose over four hours growth (see Table 1). This could explain the better adherence of B11221, and the difference in adherence between the carbon sources for B11221. Together these data suggest a correlation between the metabolism of alternative sugars and virulence phenotypes of aggregation and adhesion.

**Figure 4.**
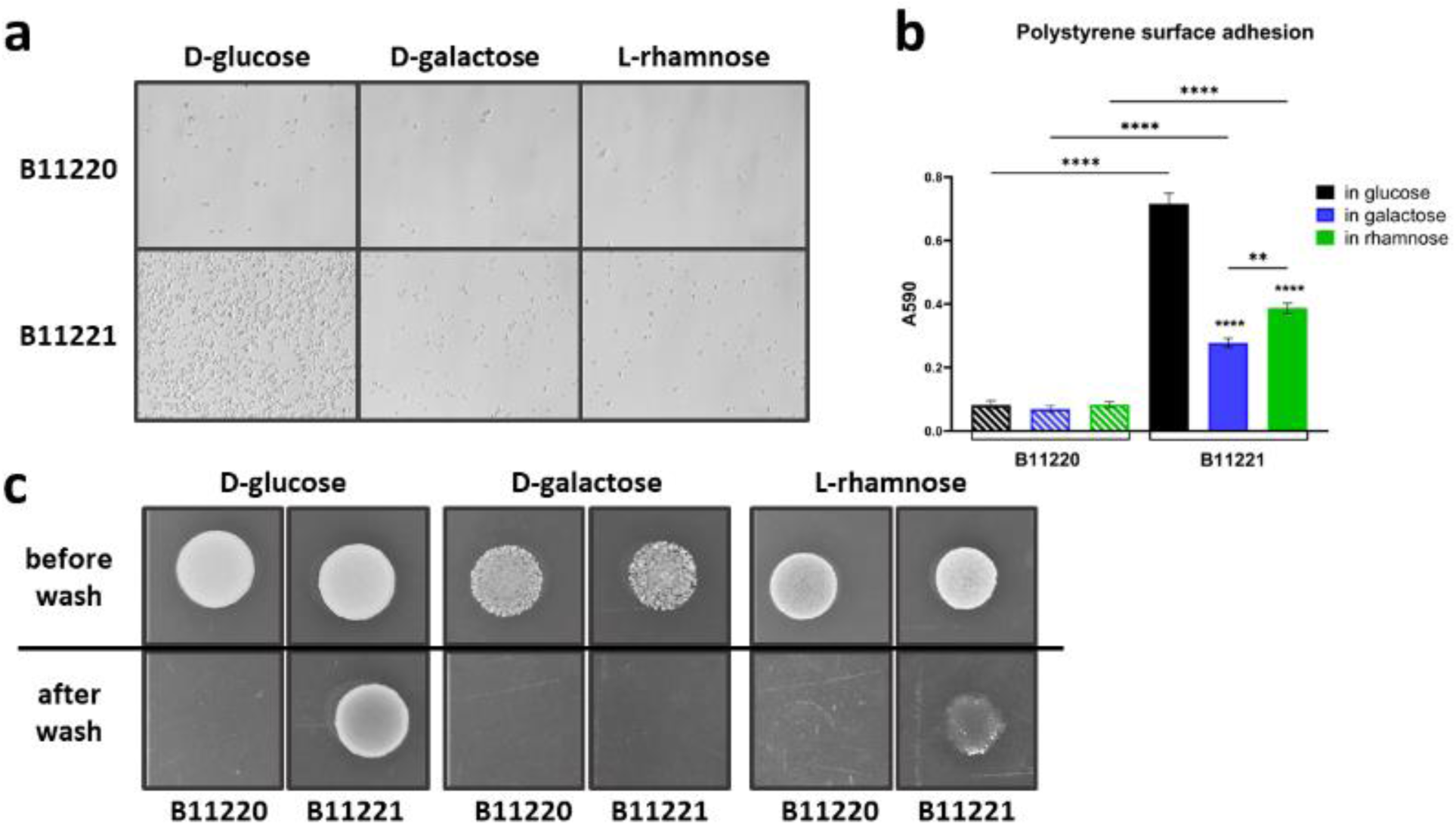
Surface adhesion characterization of B11220^Δ^ and B11221 in three carbon sources. (a) Microscopy of two *C. auris* strains cells adhered to polystyrene plate bottom after washing off the liquid culture with three sugar sources. 400x magnification. (b) Quantification of two *C. auris* strains cells adhered to the polystyrene plate bottom after 4 hours liquid culture with three sugar sources. Error bars represent the standard error (SEM). Tukey’s multiple comparisons test (****p<0.0001, **p<0.005). (c) Residue of two *C. auris* strains colony patches before and after washing off from SC agar with three sugar sources. The data shown reflects three independent replicates.

**Table 1.** ORFs unique to C. auris B11221 and their upregulation in two sugar sources

| ORFs | Function | In D-glucose |  | In Galactose |  |
| --- | --- | --- | --- | --- | --- |
|  |  | log<sub>2</sub> FC | q-value | log<sub>2</sub> FC | q-value |
| **RTCAU00203** | formamidase | 2.84 | 6.16e-51 | 1.92 | 1.01e-23 |
| **RTCAU00295** | cell-wall agglutinin | 3.28 | 4.45e-18 | ns |  |
| **RTCAU38886**<br>**RTCAU37873** | hyphal-regulated cell-wall protein /<br>Exo-alpha sialidase | ns |  | ns |  |
| **RCAUT00202** | multidrug resistance ABC transporter ATP-binding /<br>permease protein | ns |  | ns |  |
| **RTCAU20241**<br>**RTCAU28963**<br>**RTCAU38895** | GAL4 transcription factor | ns |  | ns |  |

FC: log<sub>2</sub> fold change; q-value: FDR adjusted p-value; ns: non-significant

### 3.4 Differentially expressed genes in *C. auris* B11220^Δ^ and B11221 with D-galactose

The reduced adhesion and increased tolerance to caspofungin of *C. auris* strain B11221 as compared to B11220^Δ^ when grown in D-galactose suggests metabolic alteration in B11221 during D-galactose metabolism. Thus, we analyzed the gene expression profiles of the two strains grown for four hours in aqueous SC media with 2% D-galactose or 2% D-glucose. The number of differentially expressed genes (DEGs) were greater with D-glucose than with D-galactose (952 vs. 856) in B11220^Δ^ (Fig. 5a, left). However, in strain B11221 many more genes were differentially expressed in presence of D-galactose as compared to growth in the presence of D-glucose (500 vs. 328) (Fig. 5a, right). Of these DEGs, 493 were unique to B11220^Δ^ while in the presence of D-galactose, compared to 139 in B11221 (Figure S4A). When comparing B11220^Δ^ and B11221 growing respectively in D-glucose (600 vs. 560) or D-galactose (665 vs. 775), greater numbers of genes were found to be upregulated in B11221 in presence of D-galactose (Fig. 5b), suggesting that a greater proportion of the transcriptome was altered in *C. auris* B11221 in D-galactose as compared to *C. auris* strain B11220^Δ^.

**Figure 5.**
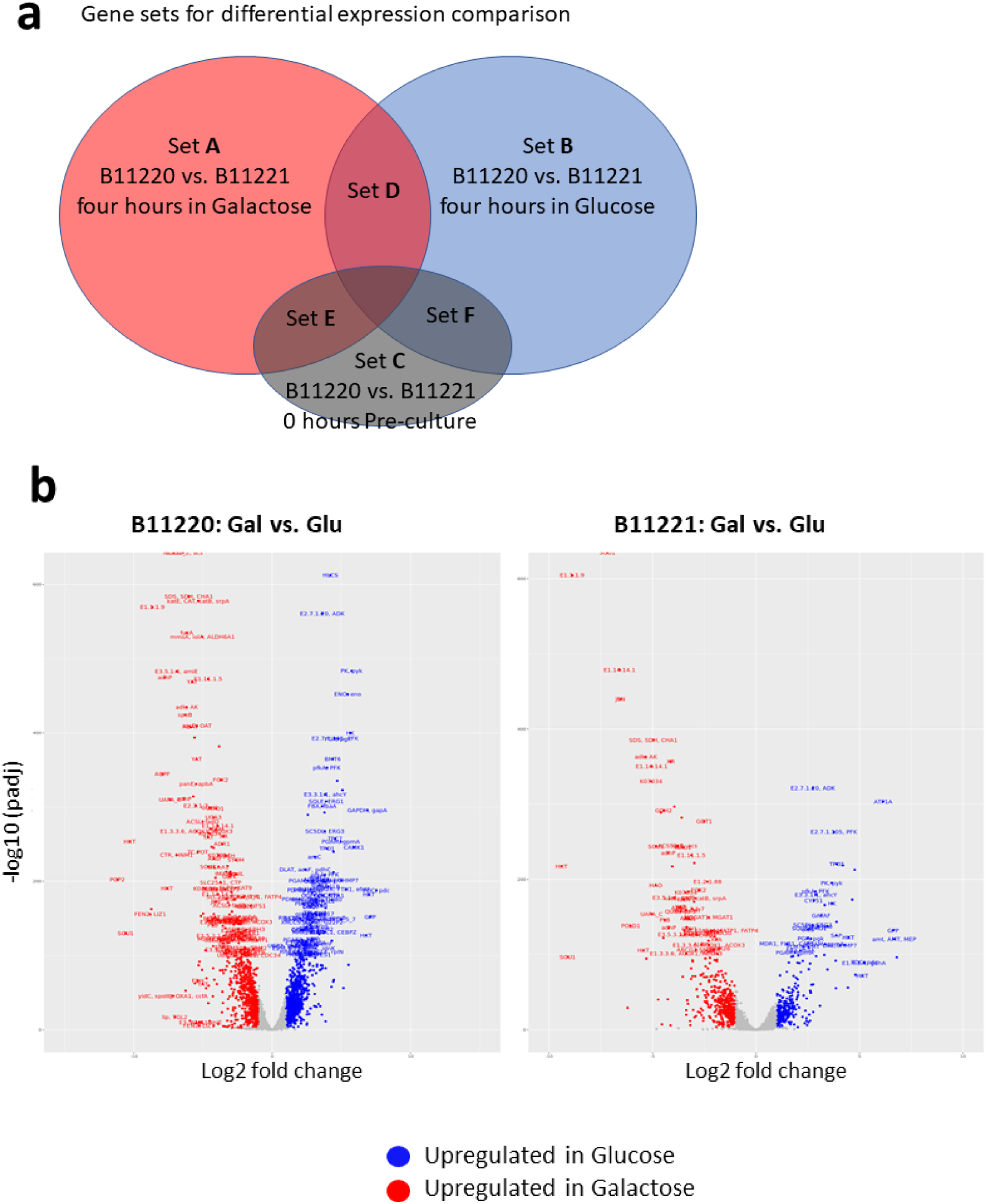
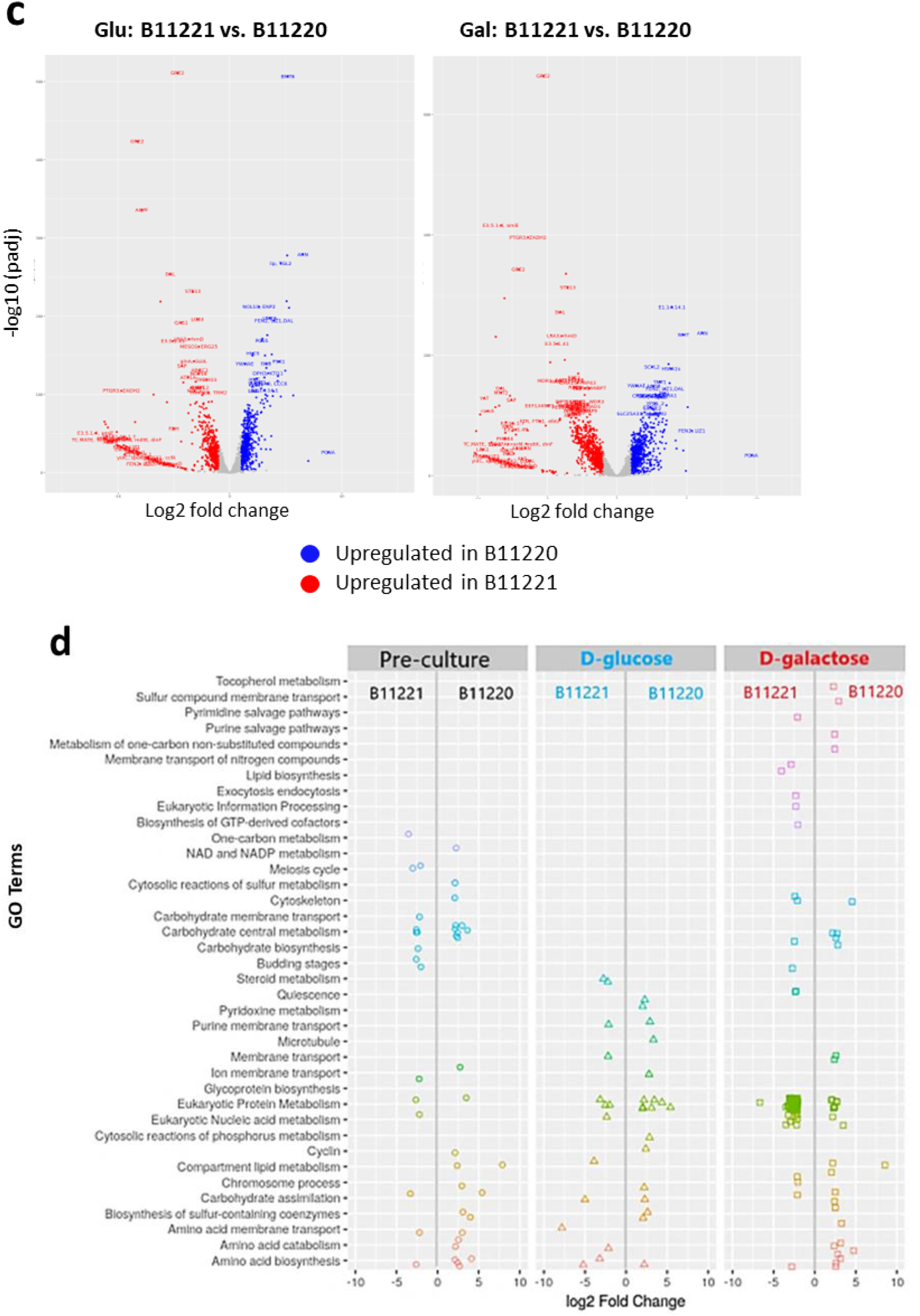
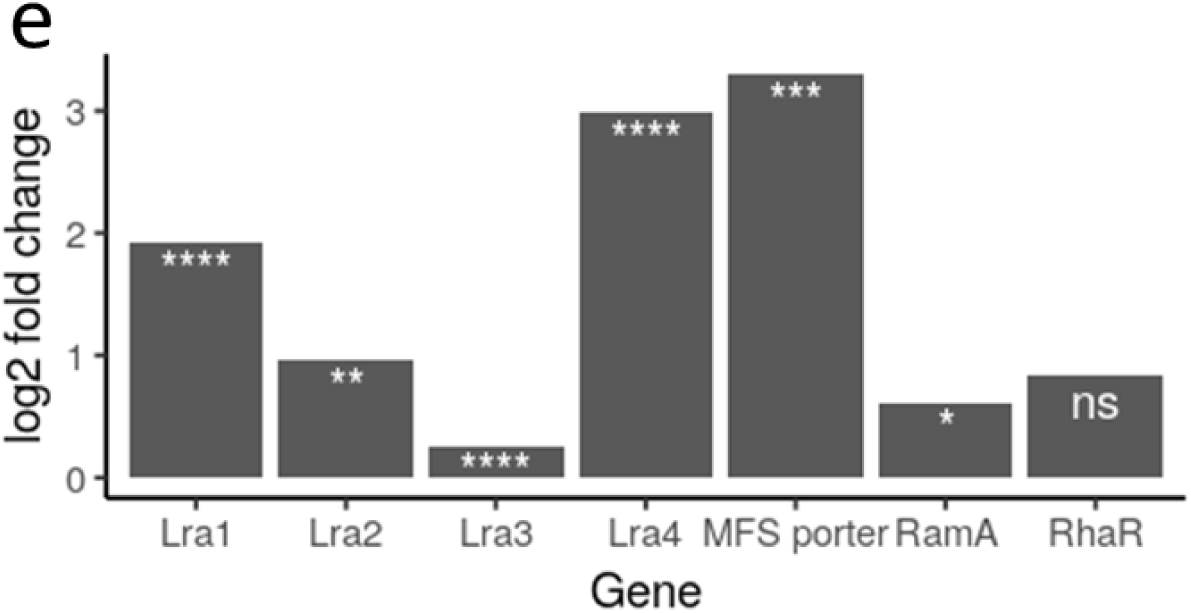
Transcriptome analysis of two *C. auris* strains in carbon source of D-glucose and D-galactose. (a) Gene sets chosen for differential expression comparison. Genes from Set A and Set B were compared for volcano plots, Set ABC were used for GO analysis, Set DEF were excluded. (b) RNA-seq volcano plots of log2 fold changes in B11220^Δ^ (left) and in B11221 (right) in presence of D-glucose (Glu) compared with D-galactose (Gal). Genes were differentially expressed (color coded) if with a fold change > 1 and an FDR adjusted p value < 0.05. Genes with a positive log2 fold change are over-expressed in D-glucose and under-expressed in D-galactose. Genes that have a negative log2 fold change are over-expressed in D-galactose and under-expressed in D-glucose. (c) RNA-seq volcano plots of log2 fold changes in B11220^Δ^ compared with B11221 respectively in D-glucose (Glu, left) and in D-galactose (Gal, right). Genes were differentially expressed (color coded) if with a fold change > 1 and an FDR adjusted p value < 0.05. Genes with a positive log2 fold change are over-expressed in B11220^Δ^ and under-expressed in B11221. Genes that have a negative log2 fold change are over-expressed in B11221 and under-expressed in B11220^Δ^. padj = adjusted p-value obtained from DESeq2. (d) GO Categorization of B11220^Δ^ comparing to B11221. Genes with a positive log2 fold change are over-expressed in B11220^Δ^ and under-expressed in B11221. Genes that have a negative log2 fold change are over-expressed in B11221 and under-expressed in B11220^Δ^. (e) Expression of seven L-rha ORFs in B11221 in presence of galactose. Y-axis represents Log2fold change. Asterisks denote False Discovery Rate adjusted p-values (q-value). **** q ≤1E−9, *** q ≤1E−4, ** q ≤ 1E−3, * q ≤ 0.005, ns not significant.

Further, we examined the gene ontology (GO) categories of the differentially expressed genes. The genes upregulated in B11221 were enriched in pathways associated with translation factors (Fig. 5c), protein metabolism (Fig. 5c), and the ribosomal complex (Fig. S3). The genes upregulated in B11220^Δ^ were enriched in genes associated with transport (Fig. S3). The results suggest that B11221 increases translation and protein production during D-galactose digestion.

All seven genes of the putative L-rhamnose gene cluster were significantly upregulated in B11221 on D-galactose(Fig. 5d). The genes encoding a MFS transporter and *LRA4* (LkaA-2-keto-3-deoxy-L-rhamnonate aldolase, EC 4.1.2.53) increased expression about three fold. *LRA1* (LraA-L-rhamnose 1-dehydrogenase, EC 1.1.1.173, 377, 378) increased one fold. ORFs of *LRA2* (LrlA-l-rhamnono-γ-lactonase, EC 3.1.1.65), *LRA3* (LrdA L-rhamnonatedehydratase, EC 4.2.1.90) and *RamA* (rgxB-α-L-rhamnosidase, EC 3.2.1.40) were also upregulated significantly in D-galactose. The upregulation of L-rhamnose degradation genes in the presence of D-galactose implies this gene cluster is functionally involved in D-galactose assimilation in *C. auris* B11221. This finding confirms that these genes are promiscuous and involved in alternate sugar metabolism rather than reserved exclusively for L-rhamnose.

The expression of some B11221 ORFs were also studied by RNA sequencing in presence of D-glucose and D-galactose (Table 1). An ORF (RTCAU00295) for cell-wall agglutinin was unique to B11221. It was upregulated in D-glucose (3.28 log2 fold-change with q-value of 4.45e-18) but not in D-galactose over four hours growth. The over-expression of agglutinin in B11221explains its aggregative phenotype when grown in liquid medium with D-glucose (Fig. 2c) and the aggregation disappearance in D-galactose. It may also contribute to the higher surface adhesion in D-glucose as well.

An ORF (RTCAU00203) for formamidase (EC 3.5.1.49) was identified in B11221 but not in B11220^Δ^. The gene for formamidase is found in other human fungal pathogens including *Paracoccidioides brasiliensis* as well as in *Helicobacter pylori*. The functional role is not very clear in pathogenesis but is involved in the hydrolysis of formamide to produce formate and toxic ammonia gas. Formate can further be converted to oxalate to feed into TCA cycle. Because this enzyme is found on the surface of hyphal cells. Previous studies have confirmed that sera of patients with proven paracoccidioidomycosis recognize the protein [85]. This gene was over-expressed in both D-glucose media (2.84 log2 fold-change with FDR-adjusted p-value of 6.16e-51) and D-galactose media (1.92 log2 fold-change with FDR-adjusted p-value of 1.01e-23).

Several ORFs unique to B11221 and located outside the 12.8 kb cluster were identified during genome sequencing and functionally categorized in Figure 6. We found four ORFs (RTCAU20241, RTCAU38895, RTCAU28963) in the *GAL4* transcription factor family, two ORFs (RTCAU38886, RTCAU37873) for hyphal-regulated cell-wall protein/exo-alpha sialidase (EC 3.2.1.18) and an ORF (RCAUT00202) for multidrug resistance ABC transporter ATP-binding/permease protein. These results, together with the biosynthetic gene cluster, point to increased galactose gene regulation in *C. auris* B11221, and the upregulated genes encode carbon source utilization, cell wall remodeling, and drug resistance genes. Therefore, alternative sugar metabolism and virulence traits may be interrelated in *C. auris*.

**Figure 6.**
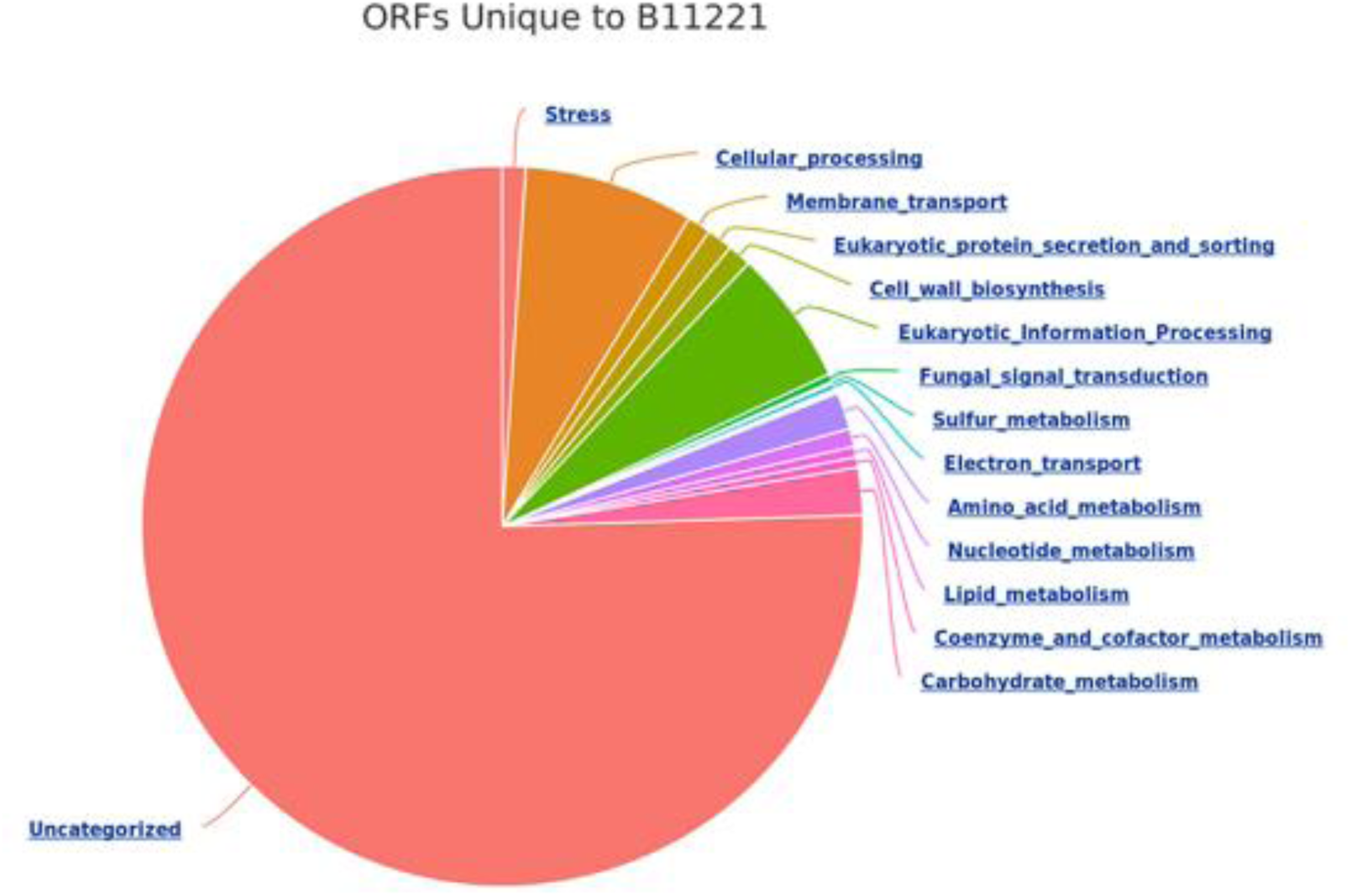
ORFs unique to B11221 categorized by their molecular function.

### 3.5 B11221 survives encounters with macrophages better than B11220^Δ^ by upregulating transporters and transcription factors

Since we noted that cell wall genes were upregulated in galactose, and cell wall composition affects immune system evasion, we next investigated the interaction of the two *C. auris* strains B11220^Δ^ and B11221 with macrophages. In mammals, macrophages are the first line of defense against microbial pathogens. Gross microscopy revealed no obvious difference between the two strains incubated with macrophages for four hours (Fig. 7a, top panels). After eight hours, more B11221 cells were unengulfed than B11220^Δ^ (Fig. 7a, bottom panels). This suggests B11221 evades macrophages better than B11220^Δ^. We then quantified macrophage survival by measuring *C. auris*colony forming units (CFU) over time. Survival of both strains was reduced after eight hours, but B11221 survival was significantly higher compared to B11220^Δ^ (Fig. 7b), which is consistent with the qualitative microscopy.

**Figure 7.**
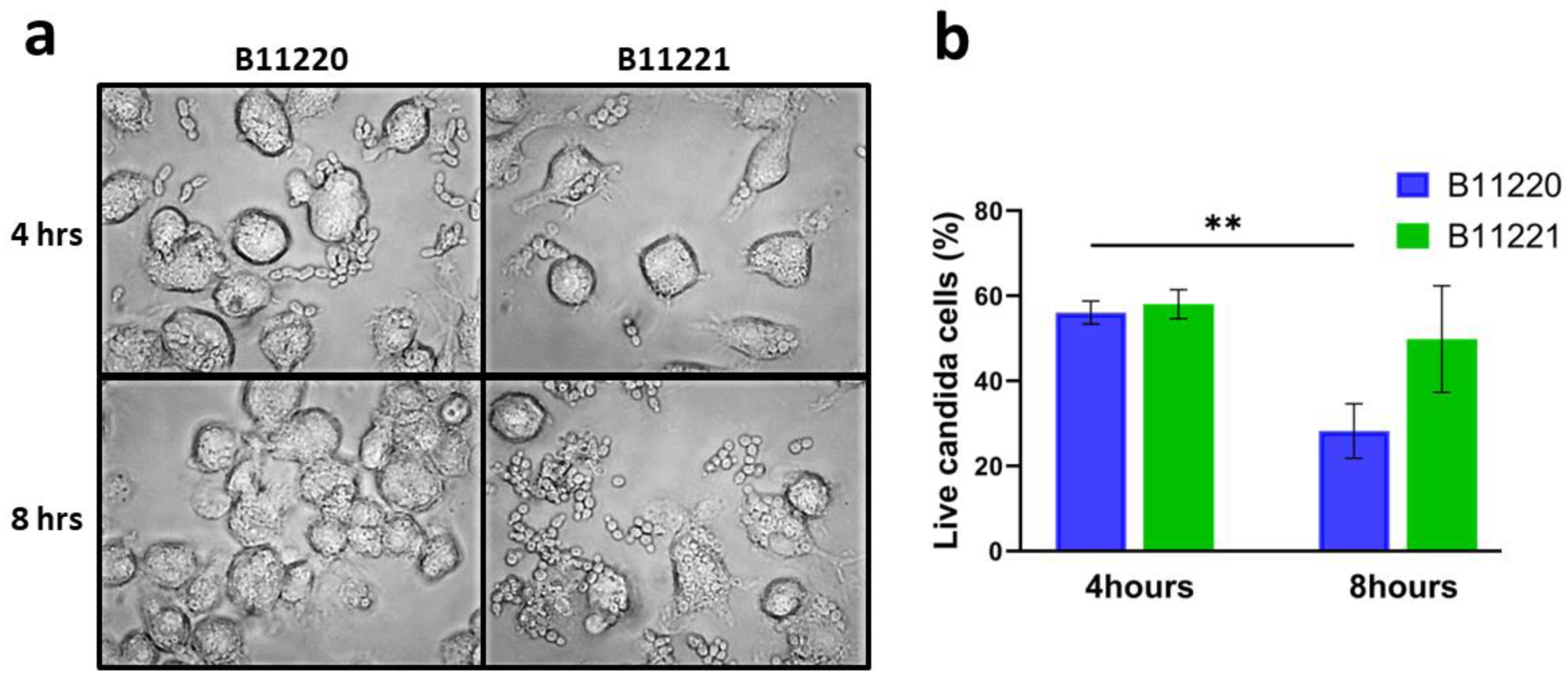
Interaction of two *C. auris* strains with macrophages. (a) Microscopy of B11220^Δ^ (left panels) and B11221 (right panels) incubation with macrophages for 4 hours (top panels) and 8 hours (bottom panels). Non-internalized Candida cells are smaller oval shaped cells around macrophages, which are larger irregular shaped cells in photos. Magnification 400x. (b) Candida survival quantification of two strains interaction with macrophages for 4 hours and 8 hours, respectively. Bars shown as mean ± SEM. Tukey’s multiple comparisons test (**p<0.005).

RNA sequencing analysis was then performed on B11220^Δ^ and B11221 when exposed to macrophages. Volcano plots of DEGs of the two strains revealed similar transcriptome patterns between two strains in conditions both without (561 vs. 438) and with (259 vs. 275) macrophage exposure (Fig. 8a). When exposed to macrophages for four hours, B11220^Δ^ showed more upregulated genes than without macrophages (113 vs. 35), and B11221 showed the same pattern (460 vs. 263) (Fig. 8b). There were 357 unique genes significantly upregulated during macrophage exposure in B11221 and only 11 unique genes in B11220^Δ^. When comparing across strains, B11221 had 124 unique upregulated genes, and only 82 in B11220^Δ^ (Figures S4C and S4D). These data suggest that both *C. auris* strains upregulate different genes in encounter with macrophages, and B11221 activates more genes.

**Figure 8.**
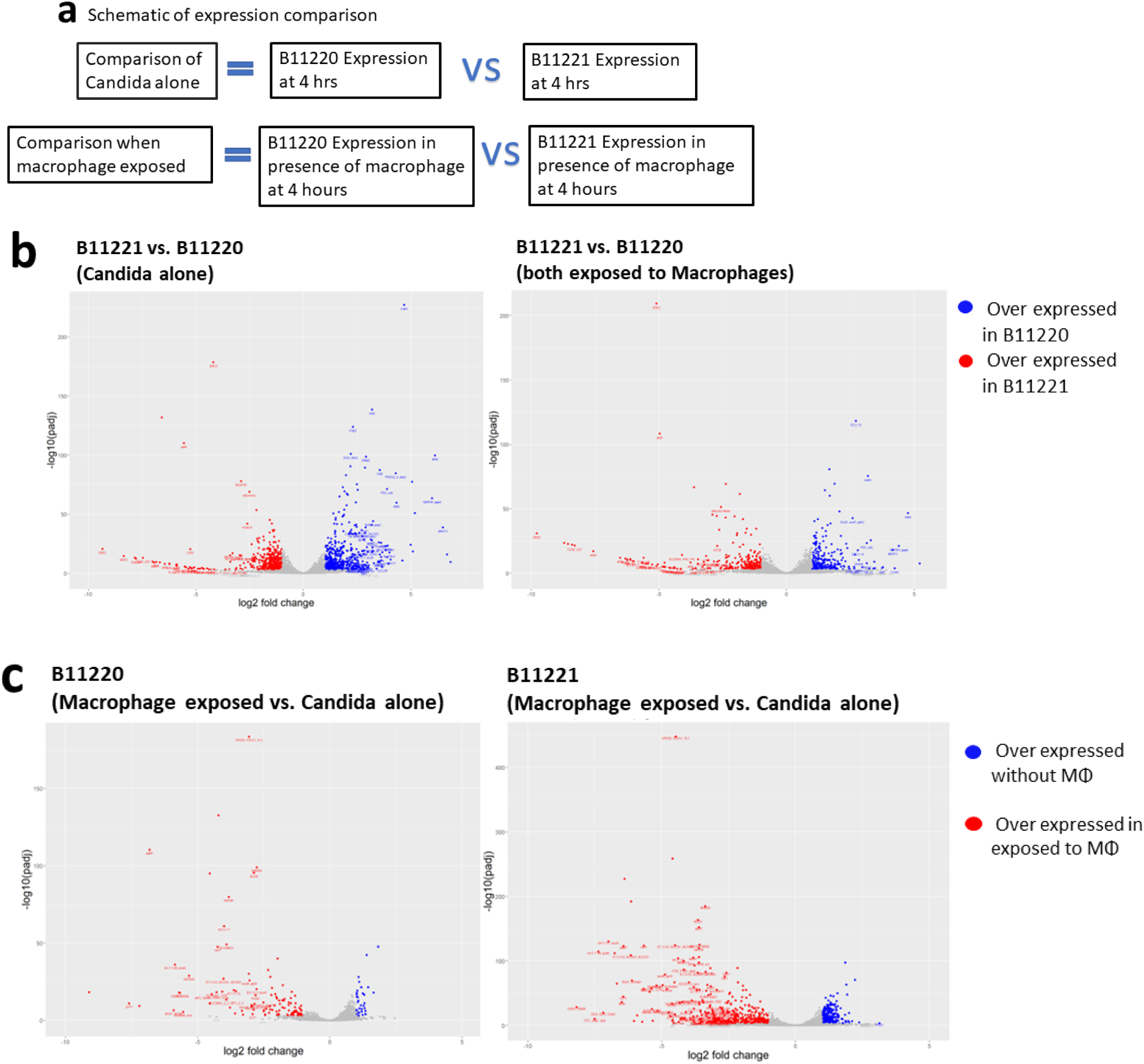
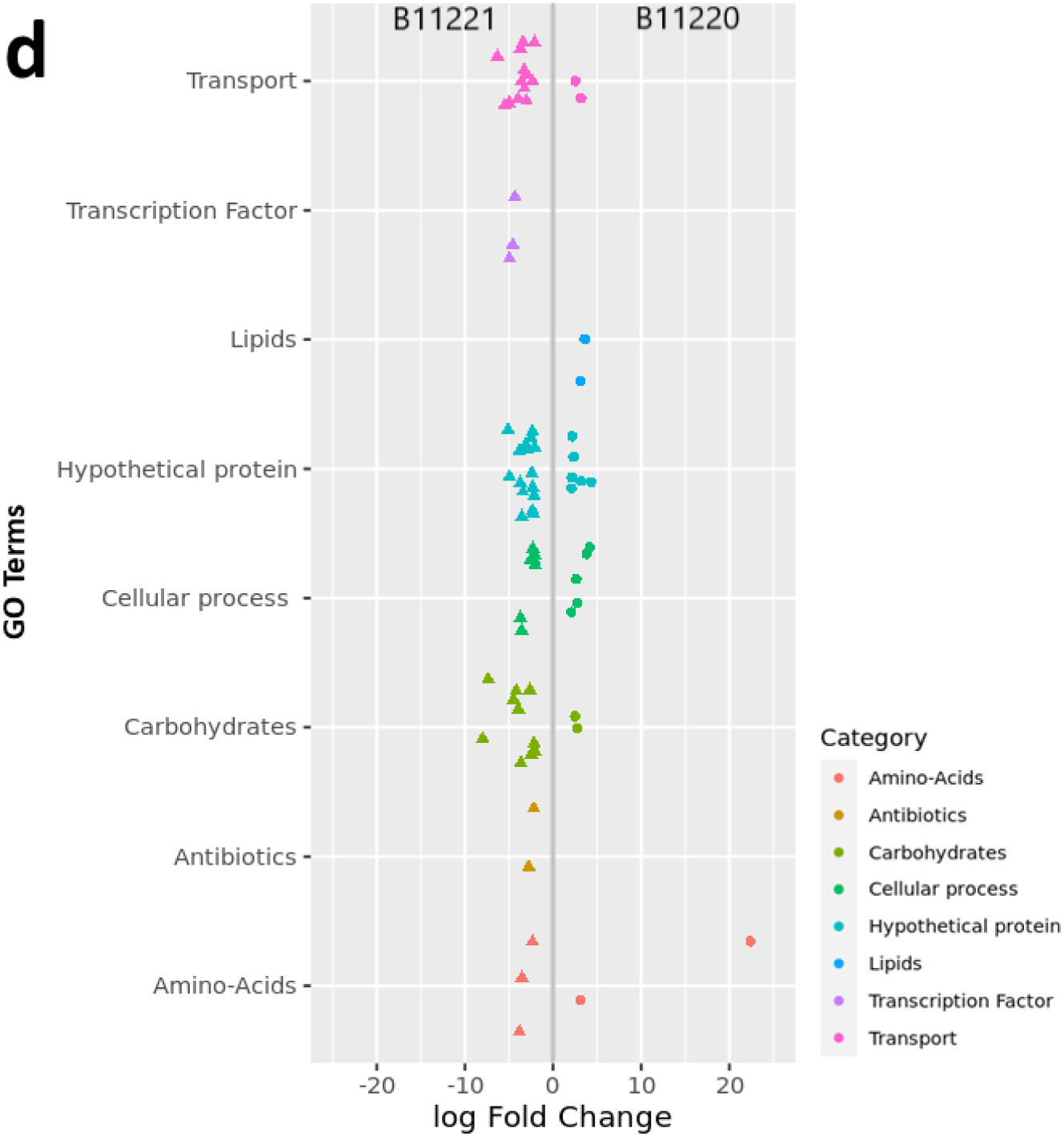
Transcriptomic analysis of interaction between macrophage and the two *C. auris* strains for 4 hours. (a) Schematic of expression comparison for RNA sequencing. For candida alone comparison, expression of two strains was compared at 4 hours culture without macrophage exposure; For comparison in presence of macrophage, expression of two strains was compared after exposed to macrophage for 4 hours. (b) RNA-seq volcano plots of log2 fold changes in B11220^Δ^ and in B11221 without (left) and in presence (right) of macrophages (MΦ). Genes were differentially expressed (color coded) if with a fold change > 1 and an FDR adjusted p value < 0.05. Genes with a positive log2 fold change (blue) are over-expressed in B11220^Δ^ and under-expressed in B11221. Genes that have a negative log2 fold change (red) are over-expressed in B11221 and under-expressed in B11220^Δ^. (c) RNA-seq volcano plots of log2 fold changes in B11220^Δ^ (left) and in B11221 (right) exposed to MΦ. Genes were differentially expressed (color coded) if with a fold change > 1 and an FDR adjusted p value < 0.05. Genes with a positive log2 fold change (blue) are over-expressed without MΦ, genes that have a negative log2 fold change (red) are over-expressed in presence of MΦ. (d) GO enrichment of genes differentially expressed in presence of macrophages but not differentially expressed without macrophage. Genes were differentially expressed if with a fold change ≥ 2 and an FDR adjusted p value < 0.05. Genes with a positive log2 fold change are up-regulated in B11220^Δ^ and down-regulated in B11221. Genes with a negative log2 fold change are up-regulated in B11221 and down-regulated in B11220^Δ^.

More genes were upregulated in B11221 in transport and transcription factors at higher levels than B11220^Δ^ (Fig. 8c, left), including *GAL4*-like transcription factors and sugar transporters (Fig. 8b), suggesting the transcription and transport activities were upregulated in B11221 during encounter with macrophages.

We then investigated whether genes already implicated in immune system evasion were upregulated (Table 2). GliC is a gliotoxin biosynthesis protein first identified in *Aspergillus fumigatus* [86]. Gliotoxin is an immunosuppressant mycotoxin that enables *A. fumigatus* to escape macrophage [87]. It is part of a larger family of genes. Previous genomics work has identified an ORF (RTCAU28975) coding for a Cytochrome P450 monooxygenase in the GliC-like family [88]. We identified proteins similar to gliP, gliG, gliI, gliT, and gliN in both B11220^Δ^ and B11221. However, proteins similar to gliK and gliJ were not found. These genes were not found to be significantly differentially expressed when exposed to macrophages. We also found two genes (RTCAU08811, RTCAU21082) that are identified as NADPH-dependent methylglyoxal reductase (EC 1.1.1.283), which is part of methylglyoxal degradation. These are significantly over-expressed in B11221 compared to B11220^Δ^ (5.10 and 9.79 log2 fold-change, with FDR-adjust p-value of 1.22e-91 and 3.51e14), but not significantly overexpressed in B11221 when in presence of macrophage. However, we found that an ORF encoding Candidapepsin (RTCAU28889), a virulence factor that degrades host proteins [89], was overexpressed in B11221 (4.15 log2 fold-change, FDR adjusted p-value 0.0492).

**Table 2.**
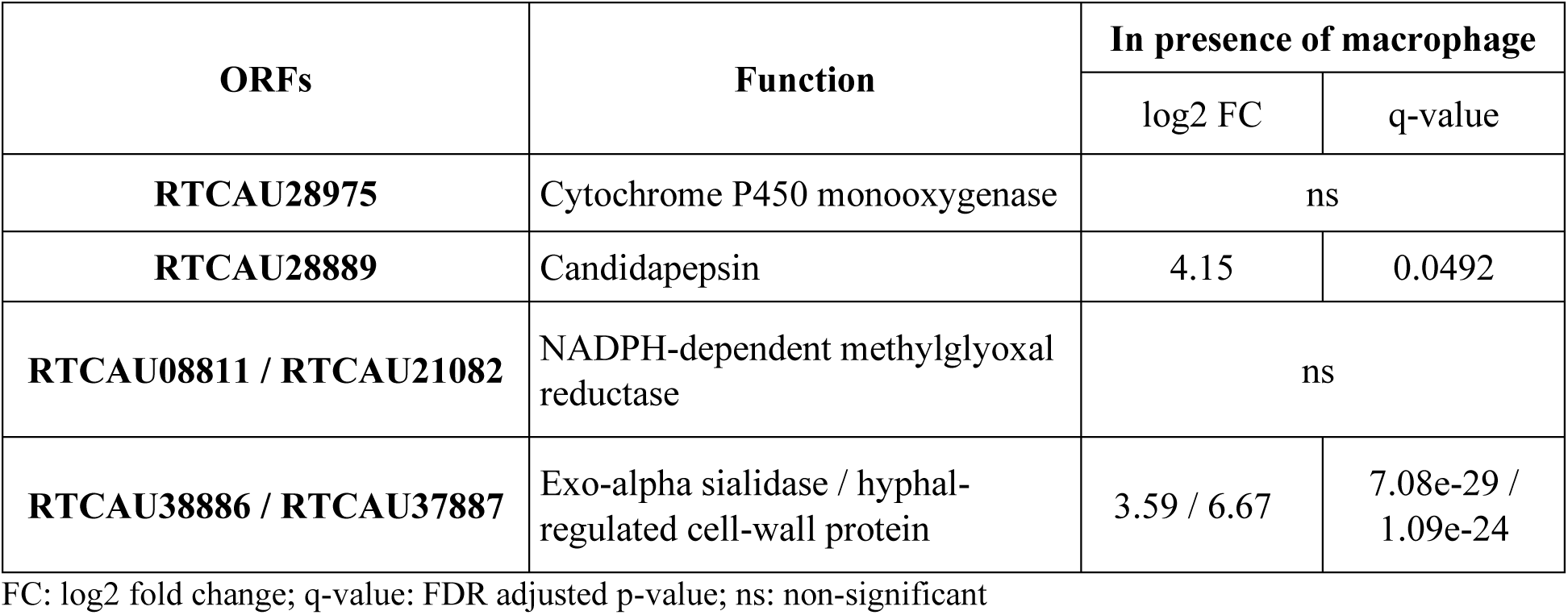
ORFs unique to C. auris B11221 and their upregulation in presence of macrophage

| ORFs | Function | In presence of macrophage |  |
| --- | --- | --- | --- |
|  |  | log2 FC | q-value |
| **RTCAU28975** | Cytochrome P450 monooxygenase | ns |  |
| **RTCAU28889** | Candidapepsin | 4.15 | 0.0492 |
| **RTCAU08811 / RTCAU21082** | NADPH-dependent methylglyoxal reductase | ns |  |
| **RTCAU38886 / RTCAU37887** | Exo-alpha sialidase / hyphal-regulated cell-wall protein | 3.59 / 6.67 | 7.08e-29 / 1.09e-24 |

FC: log2 fold change; q-value: FDR adjusted p-value; ns: non-significant

Finally, two of the most upregulated genes in B11221 were found to be RTCAU38886 (8.56 FDR adjusted p-value of 1.36e-10) and RTCAU37887 (log2 fold-change 8.71 with FDR-adjusted p-value of 5.78e-11). They were identified as Exo-alpha sialidase (EC 3.2.1.18) / Hyphally regulated cell wall protein. This protein contains a domain similar to that of *HYR1* in *C. albicans*. *HYR1* is a macrophage-induced cell-wall protein [90] with a role in innate immune cell evasion [91]. Both genes are also significantly overexpressed when B11221 is in the presence of the macrophage (3.59 and 6.67 log2 fold change, FDR-adjusted p values of 7.08e-29 and 1.09e-24). Taken together, these results show that *C. auris* B11221 is more responsive to macrophage exposure, and that response activates candidapepsin and a HYR1-like gene that could mediate improved macrophage survival.

### 3.6 B11221 has lower β-glucan on the cell surface compared to B11220^Δ^

Since several genes involving the cell wall were implicated in our carbon source study, and cell wall proteins were implicated in our macrophage study, we characterized the β-(1,3)-glucan exposure in the cell wall of *C. auris* B11220^Δ^ and B11221. β-glucan masking has been shown to affect fungi immune responses [56, 92] and β-(1,3)-glucans make up approximately 40% of the *Candida* cell wall [55].

We compared the surface exposure of β-(1,3)-glucan between the two strains using flow cytometry and found more β-(1,3)-glucans are exposed on the cell surface of B11220^Δ^ as compared to *C. auris* strain B11221 (Fig. 9a). β-glucan masking has been shown to affect fungi immune responses [56, 92] and β-(1,3)-glucans make up approximately 40% of the *Candida* cell wall [55]. The differences in β-(1,3)-glucan exposure between the two strains may contribute to their phenotypic variations. To eliminate the interference of other cell wall components masking the glucans, we also measured total β-(1,3)-glucan present on the surface of the two strains using Enzyme Linked Immune Sorbent Assay (ELISA). Our results indicate that stain B11220^Δ^ contains more β-(1,3)-glucan (Fig. 9b), These results agree with our observation that B11221 is more resistant to caspofungin than B11220^Δ^ (Fig. 3). The lower content of β-(1,3)-glucan required on B11221 to maintain its normal function may enable it to be less affected by caspofungin which targets the biosynthesis of β-(1,3)-glucan. Furthermore, limited β-(1,3)-glucan exposure on B11221 surface likely leads to decreased recognition by macrophages allowing them to avoid phagocytosis (Fig. 7a, bottom panels) and better survival upon an encounter with macrophages as compared to B11220^Δ^ (Fig. 7b). Together, these results provide strong evidence that metabolism and the cell wall are linked to antifungal resistance and immune system evasion in *C. auris* B11221.

**Figure 9.**
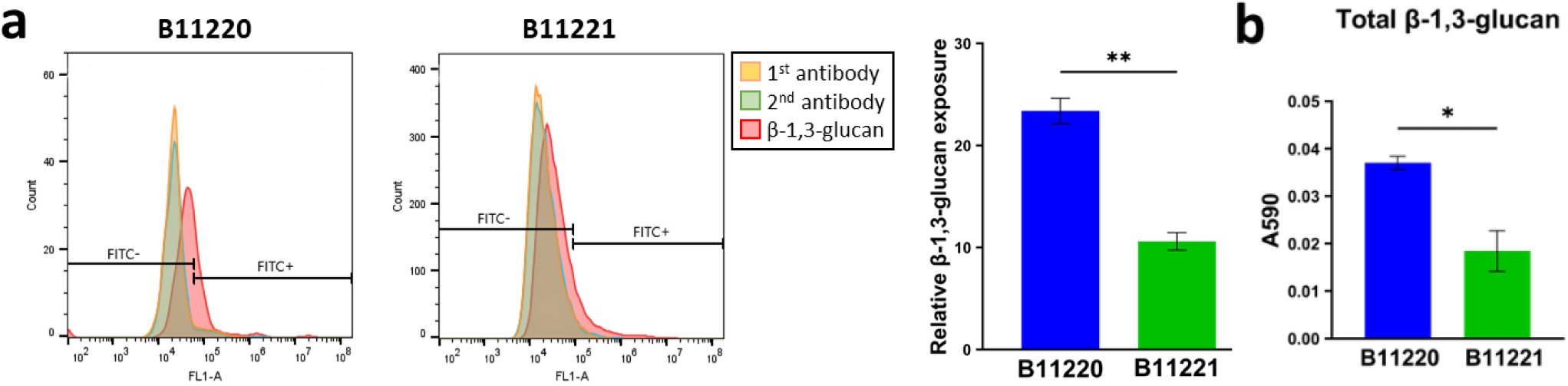
Quantitative analysis of cell wall β-1,3-glucan in two *C. auris* strains. (a) Representative flow cytometry analysis of β-1,3-glucan exposure on *C. auris* cell wall. Yellow histograms correspond to control with only anti-β-glucan MAb, green histograms correspond to control with only anti-Mouse IgG Secondary Antibody, and the red histogram represents sample with both antibodies. Red peak shifts on the plots from left to right indicate elevated fluorescence intensities indicating increasement in the β-1,3-glucan exposure. Quantitated results are shown on the right. (b) The total cell wall β-1,3-glucan content on B11220^Δ^ and B11221. Bars shown as mean ± SEM. Unpaired t test (*p<0.05, **p<0.005).

## 4 Conclusion and Discussion

Since its emergence as a multi-drug resistant fungal pathogen in 2009, over 12 genomes of *Candida auris* have been reported so far [93]. Here, we used both short read (Illumina) and long read (Nanopore) sequencing techniques to ensure a more continuous sequence result of two species of *C. auris* that represent the two ends of drug susceptibility, strain B11220^Δ^ (Clade II) and B11221 (Clade III). Using these improved genomes, we confirmed a large deletion that encodes a cluster of seven genes involved in the metabolism of alternate sugars. This cluster of genes is absent in Clades I, II, IV, and present in Clade III and Clade V [22]. Clade III isolates are reported to be able to assimilate L-rhamnose, but not Clades I, II, and IV isolates [94], which may be because the gene cluster is missing in these clades. The Clade V isolate also conserved the gene cluster like Clade III isolates, supporting the hypothesis that in *C. auris* the absence of this cluster in Clades I, II, and IV is a loss rather than a gain in Clade III [22]. High rate of genome rearrangements and subtelomeric loss are also features reported unique to Clade II isolates [22], underlying their phenotype of being more sensitive to UV-C killing [95, 96] and drug susceptible. However, they are still potentially able to cause severe, drug-resistant infections as some isolates have been reported to acquire azole resistance [97]. The fact that *C. auris* B11221 (Clade III) contains the gene cluster and is able to digest L-rhamnose is rare [98, 99] [100], and the loss of the L-rha gene cluster in B11220^Δ^ may connect to the geological origin changes and hosts colonized.

In addition to the 12.8 kb deletion, our whole genome sequencing identified several ORFs that were unique to B11221. These ORFs could contribute to the phenotypic differences between B11221 and B11220^Δ^, including agglutinin (RTCAU00295) which is involved in aggregation and adhesion [101], a component of the multidrug resistance ABC transporter (RCAUT00202) [102], a hyphal-regulated cell wall protein implicated in virulence (RTCAU38886) [103], a sialidase (RTCAU37873) associated with macrophage survival and alternative sugar utilization [104], and formamidase (RTCAU00203) which is found in other human fungal pathogens but the role is not fully understood [100].

We also looked at glucan synthase mutations, since they are responsible for resistance to echinocandin drugs (caspofungin) [36]. We did not identify the resistance-associated hot spot region mutations in *FKS1* [40] in the two strains. However, new *FKS* mutations related to drug resistance have been described [105] and a more sensitive and robust method has been developed [106], which can be applied in *C. auris* to detect either new mutations or non-*FKS1*-linked echinocandin resistance mechanisms. Overall, our genome assemblies point to several key genes in *C. auris* B11221 that may mediate its increased virulence.

Our transcriptomic studies also identified key rewiring and upregulation of genes in B11221 when grown on alternative sugars and when exposed to macrophages. We report a larger transcriptomic change in B11221 (Clade III) when metabolizing D-galactose as compared to B11220^Δ^ (Clade II). The activated genes are involved in galactose metabolism as well as cell wall composition and antifungal resistance. The galactose grown B11221 cultures lost aggregative form in liquid medium, decreased adherence to abiotic surface, became more drug susceptible, yet developed drug tolerance to caspofungin. The aggregation trait is linked to transcriptional changes in genes associated in surface adhesion in *C. auris* [107, 108], with potential clinical implications [83].

Previous studies on *C. albicans* reported that growth in galactose altered the outermost surface components of the fungal cell wall [109, 110], and consequently increase the synthesis of outer fibrillar-floccular layer mannoprotein adhesins [111, 112] that facilitate fungal adhesion and biofilm formation [112, 113], thus the adhesion increases relative to growth in glucose [110, 114]. In contrast, our results revealed decreased adhesion of B11221 in D-galactose and L-rhamnose, suggesting the metabolism of carbon source can be very different from *C. albicans* to *C. auris*, especially the strains with the sugar metabolism gene cluster. Importantly, *C. auris* has an affinity for skin, unlike other *Candida* species tending to colonize the gut. This increases the chances for *C. auris* to spread within and between healthcare networks when colonized or infected patients are transferred. Thus, decreasing attachment may contribute to its easy spread in healthcare settings [112] [113].

We also found that *C. auris* B11221 developed caspofungin tolerance when grown the presence of glucose and galactose. This indicates that while galactose decreases adherence, it is more likely that B11221 will develop drug resistance on alternative sugars. Clearly, there is a need to investigate this observed tolerance, since cells in the tolerant state are often isogenic to those in the non-tolerant state [44].The state of the host’s innate immunity plays a major role in the establishment of infections caused by opportunistic fungal pathogens such as *Candida* spp. Our results showed that as early as 4 hours of interaction period, the two *C. auris* strains survived macrophages at a similar percentage, however, phagocytosis observations at 4 hours revealed that B11220^Δ^ cells were already internalized by macrophages, as the bridges forming and stretching shapes were observed (Fig S2, indicated by red arrow), while B11221 cells mostly only attached around the surface of the macrophages and were not internalized. Our transcriptomic analysis showed that the interaction of macrophage cells on the two strains differentiated at an earlier stage. B11221 survived phagocytosis better than B11220^Δ^ and upregulated transcription factors and genes associated in transport when encountered macrophage as compared to B11220^Δ^.

There could be two basic strategies that *Candida* survives macrophage encounters: 1) escaping from macrophage digestion so *Candida* can survive phagocytosis; or 2) reducing the recognition by macrophage receptors so they escape phagocytosis. The evidence presented that B11221 has less surface β-glucan and survives macrophages better supports the latter hypothesis. However, the first strategy cannot be completely ruled out. Even though it is not differentially expressed, the GliC-like family ORF (RTCAU28975) is expressed, making it possible that *C. auris* is synthesizing gliotoxin, an immunosuppressant mycotoxin [87]. In either case, it is clear that *C. auris* B11221 rewires its transcriptome and expresses key genes involved in macrophage evasion, and these genes are influenced by growth on D-galactose.

Therefore, this study of two *C. auris* isolates representing two distinct clades showed that *C. auris* may have evolved effective immune evasion strategies, environmental nutrient adaptation, and stress responses to ensure its host colonization, especially for *C. auris* B11221 which is in the outbreak causing clade. Thus, this study defines important genes involved in the virulence mechanisms of *C. auris*, an emerging fungal pathogen, and provides genetic evidence linking metabolism, drug resistance, and immune system evasion in pathogenic yeasts.

